# Metabolic Profiles of Encapsulated Chondrocytes Exposed to Short-Term Simulated Microgravity

**DOI:** 10.1101/2024.07.01.601604

**Authors:** Annika R. Bergstrom, Matthew G. Glimm, Eden A. Houske, Gwendolyn Cooper, Ethan Viles, Marrin Chapman, Katherine Bourekis, Hope D. Welhaven, Priyanka P. Brahmachary, Alyssa K. Hahn, Ronald K. June

**Author notes:** <u>Co-corresponding Authors</u> Ronald K. June, Ph.D. Dept. of Mechanical & Industrial Engineering Montana State University PO Box 173800, Bozeman, MT 59717-3800; ORCID: 0000-0003-0752-4109; Alyssa K. Hahn, Ph.D. Dept. of Biological & Environmental Sciences Carroll College 1601 N Benton Ave, Helena, MT, 59625; ORCID: 0000-0002-8884-2610.

## Abstract

The mechanism by which chondrocytes respond to reduced mechanical loading environments and the subsequent risk of developing osteoarthritis remains unclear. This is of particular concern for astronauts. In space the reduced joint loading forces during prolonged microgravity (10^-6^ *g*) exposure could lead to osteoarthritis (OA), compromising quality of life post-spaceflight. In this study, we encapsulated human chondrocytes in an agarose gel of similar stiffness to the pericellular matrix to mimic the cartilage microenvironment. We then exposed agarose-chondrocyte constructs to simulated microgravity (SM) using a rotating wall vessel (RWV) bioreactor to better assess the cartilage health risks associated with spaceflight. Global metabolomic profiling detected a total of 1205 metabolite features across all samples, with 497 significant metabolite features identified by ANOVA (FDR-corrected p-value < 0.05). Specific metabolic shifts detected in response to SM exposure resulted in clusters of co-regulated metabolites, as well as key metabolites identified by variable importance in projection scores. Microgravity-induced metabolic shifts in gel constructs and media were indicative of protein synthesis, energy metabolism, nucleotide metabolism, and oxidative catabolism. The microgravity associated-metabolic shifts were consistent with early osteoarthritic metabolomic profiles in human synovial fluid, which suggests that even short-term exposure to microgravity (or other reduced mechanical loading environments) may lead to the development of OA.

## Introduction

Osteoarthritis (OA) is the most common degenerative joint disease, affecting over 500 million individuals worldwide^1^. OA is a whole joint disease associated with changes to the subchondral bone (*e.g.*, osteophyte formation and subchondral bone sclerosis), inflammation of the synovial membrane, and articular cartilage (AC) degradation, the hallmark. These changes in whole joint structure lead to significant pain, joint stiffness and reduced mobility, and an overall reduced patient quality of life. Current treatments include pain management for early OA and joint replacements for end-stage disease.

A major risk factor for the development of OA is abnormal mechanical loading. Deviations from physiological loading conditions (*e.g.,* high impact forces^2, 3^ or reduced loading^4, 5^) result in a metabolic imbalance in articular chondrocytes. This imbalance favors matrix catabolism, which weakens cartilage and diminishes the ability to resist mechanical loads^6^. While other musculoskeletal tissues, such as bone and muscle can reverse the effects of abnormal loading, AC is avascular, and thus has a poor regeneration capacity^7^. This results in potentially irreversible AC degeneration associated with that of OA.

Microgravity exposure (10^-6^ *g*) significantly reduces the mechanical loads applied to the musculoskeletal system. This may have deleterious effects on the AC and increase the risk of developing OA. In studies analyzing the effects of joint immobilization after clinical bed-rest, reduced loading results in significant reductions in cartilage thickness at weight-bearing regions of the tibia at short-term time points (two weeks^4^) and at the patella, medial tibia, and medial and lateral femoral condyle at longer time points (12-24 months post-injury^8,9^). Mouse studies of reduced loading (either by spaceflight or terrestrial hind-limb unloading) found decreases in the levels of glycosaminoglycans and collagen, increased concentrations of catabolic enzymes (MMP-13), and reduced cartilage thickness^10-14^. Very few studies have investigated the effects of microgravity exposure on human cartilage and chondrocytes. Analysis of urine and serum markers of cartilage degradation found elevated levels of cartilage markers in human biofluids in the 30 days following long-duration (>5 month) microgravity exposure^15-17^. Taken together, these studies provide clear evidence that reduced mechanical loading due to microgravity leads to cartilage degeneration. However, more data are needed to fully understand the underlying mechanisms and pathways driving these degenerative changes.

Thus, the overall goal of this work is to expand our understanding of chondrocyte mechanotransduction in reduced loading environments to help uncover the underlying mechanism driving these degenerative changes to better assess astronaut risk of OA post-spaceflight. We hypothesized that exposing encapsulated chondrocyte cells to SM would induce shifts in metabolism. To address this hypothesis, human chondrocytes were encapsulated in an agarose gel of similar physiological stiffness to the pericellular matrix (PCM) to better mimic the cartilage biomechanical environment and exposed to simulated microgravity (SM) using a rotating wall vessel (RWV) bioreactor. The response to simulated microgravity was analyzed using global metabolomic profiling to gain an untargeted and unbiased view of metabolic shifts.

## Methods

### Cell culture

The human chondrosarcoma cell line SW1353 (HTB-94) was purchased from the American Type Culture Collection and cultured in a complete high glucose Dulbecco’s modified eagle’s medium (DMEM) supplemented with 1% Penicillin-Streptomycin and 10% fetal bovine serum. The cell line was incubated in a 37°C humidified atmosphere with 5% CO_2_, and cultures were maintained at < 80% confluency.

### Chondrocyte encapsulation

SW1353 cells were encapsulated using an established protocol^18,19^. Cells were encapsulated in 4.95% agarose gels at a concentration of 800,000 cells/gel. Gels were cylindrical with a height of 10.67 mm and diameter of 6.86 mm. This cell concentration was chosen based on the density of chondrocytes in human articular cartilage^20^. Briefly, agarose powder and water were combined in a beaker and heated for 10 second intervals until the mixture became homogenous. A portion of the agarose mixture was removed and SW1353 cells were added and gently mixed. The cell-agarose mixture was then transferred into wells of a 96-well plate and allowed to solidify and polymerize^19^.

### Experimental design

Samples of agarose-encapsulated chondrocytes were created and individually weighed for consistency (0.329 ± 0.006 mg). Half (n = 16) of the solidified encapsulated chondrocytes were placed into Rotary Cell Culture System (RCCS) vessels and filled with media for exposure to SM on the RWV bioreactor (Synthecon, Houston, TX) (SM cohort). Normal gravitational force controls were generated using the remaining encapsulated chondrocytes (n = 16) by placing each gel into a RCC vessel, filling with media, and incubating for exposure to normal gravity.

RCCS vessels were attached to the RWV bioreactor inside of the incubator (37°C, 5% CO_2_). The RWV bioreactor was manually set to 13.2 rpm to correctly position the encapsulated chondrocytes in constant free fall based on pilot experiments. The vessels were carefully monitored throughout the duration of the experiments to maintain constant free fall and ensure the absence of bubbles. The remaining RCCS vessels were left in the incubator (37°C, 5% CO_2_) to serve as controls. Both SM and control encapsulated chondrocytes were maintained under either 10^-6^ *g* or normal gravitational forces for a period of four days (Figure 1).

**Figure 1.**
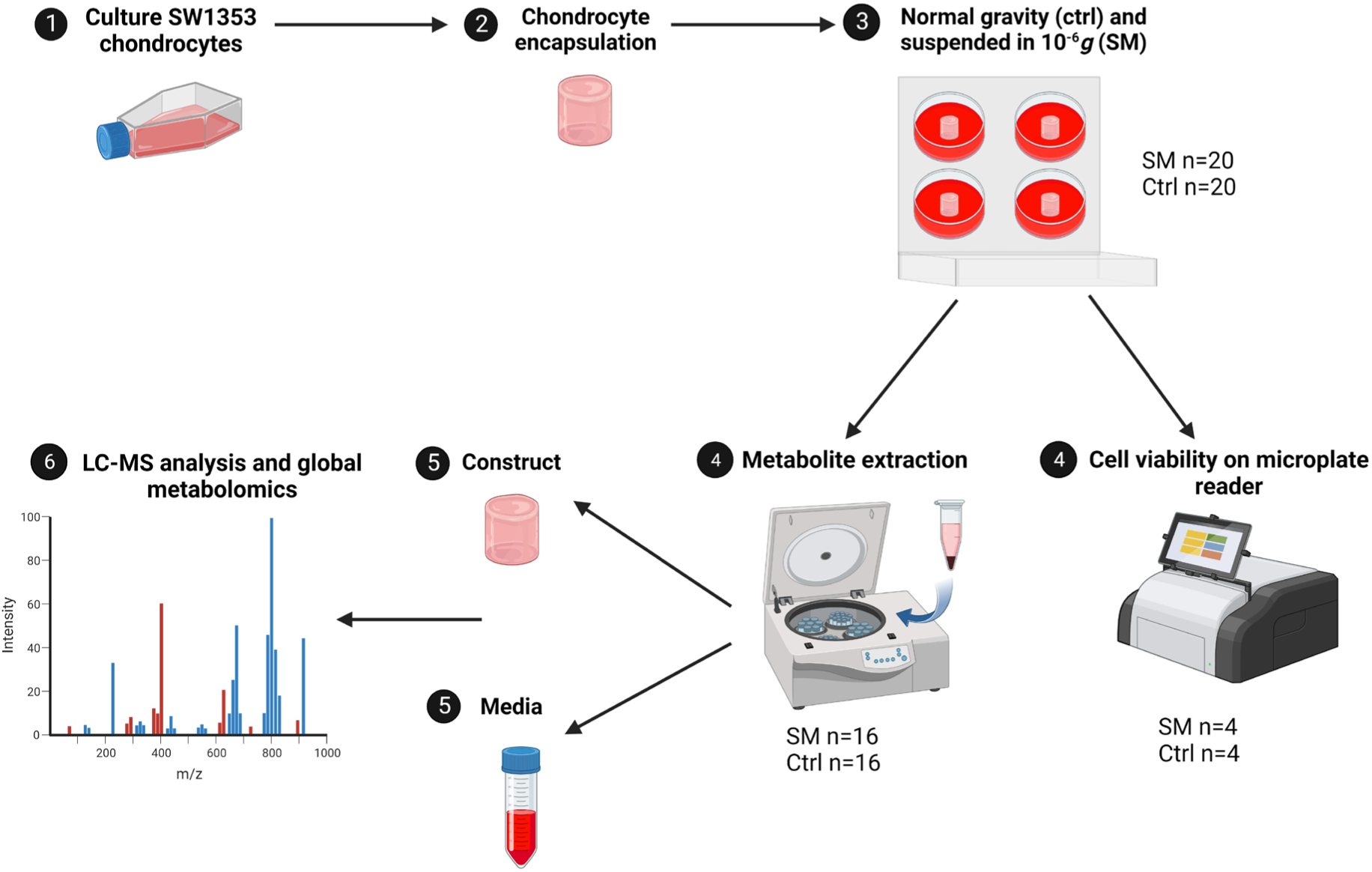
Experimental design of study. SW1353 cells were cultured to P10-14, cells were encapsulated in an agarose gel construct, and suspended in either normal gravitational forces in a CO_2_ incubator or microgravity (10^-6^*g*) on a RWV bioreactor. Cell viability was confirmed through RFUs on a microplate reader (SM n=4, Ctrl n=4). Metabolite extractions were performed on both SM (n=16) and control (n=16) groups and ran on an LC-MS for metabolomic profiling. SM = simulated microgravity and Ctrl = normal gravity.

### Cell viability

A subset of the encapsulated chondrocytes was assessed for cell viability at the end of the four-day timepoint. SM (n = 4) and control (n = 4) encapsulated chondrocytes were washed once with phosphate-buffered saline (PBS) and incubated in a solution containing 8 µmol calcein AM and 75 µmol propidium iodide for 1 hour at 37℃ before analyzing on a Thermo Fisher Scientific Varioskan LUX multimode microplate reader. Using an established protocol, relative fluorescence units (RFUs) were obtained for calcien AM (ex/em 490/517 nm) representing live cells and propidium iodide (ex/em 535/617 nm) representing dead cells^21^. The means and standard deviations were then determined.

### Metabolite extraction

Metabolites were extracted from encapsulated chondrocytes and media using an established metabolite extraction protocol, with slight modifications/adaptions for agarose gel^22^. From each vessel, 50 mL of media was harvested and centrifuged at 500 x *g* at 4°C for 5 minutes to remove debris and a second time for 10 minutes at 12,000 rpm. Encapsulated chondrocytes were flash frozen in liquid nitrogen to halt metabolism and homogenized using the SPEX SamplePrep tissue homogenizer (SPEX SamplePrep, Metuchen, NJ) with 1 mL of 3:1 methanol:water prior to centrifugation. Supernatants were collected and mixed with 80% v/v methanol:water at -20°C for 30 minutes (other 20% miliQ/RO water) to extract metabolites followed by vortexing for 3 minutes. Samples were centrifuged at 16,100 x *g* for 5 minutes at 4°C, and the supernatants were transferred to a new tube before metabolites were dried down in a vacuum concentrator (Savant AES 1010, ThermoFisher Scientific, Waltham, MA). Proteins and lipids were precipitated by re-extracting with 5 volumes (1 mL each = 5 mL total) of aqueous 1:1 (v/v) acetonitrile:water v/v at 0°C for 30 minutes. Samples were centrifuged at 16,100 x *g* a final time for 5 minutes and the supernatants were collected and dried down in the vacuum concentrator. All samples were then stored at -80°C until ready for mass spectrometry analysis. All extraction solvents were HPLC-grade or higher.

### Liquid chromatography-mass spectrometry analysis

Dried metabolite extracts were resuspended in mass spectrometry grade 50:50 water:acetonitrile as the liquid chromatography-mass spectrometry (LC-MS) injection solution. The LC-MS system used for analysis was an Agilent 1290 UPLC connected to an Agilent 6538 Quadrupole-Time of Flight (Q-TOF) mass spectrometer (Agilent, Santa Clara, CA) operated in positive mode. Separation of metabolites occurred on a Cogent Diamond Hydride HILIC column (150 x 2.1 mm, MicroSolv, Eatontown, NJ) using an optimized normal phase 25-minute gradient elution method^21^.

### Statistical analysis

Data generated by the LC-MS were initially processed using MSConvert^23^ and XCMS^24^, as previously described^22^. Metabolomic profiling was then performed using MetaboAnalyst (version 4.0)^25^. Prior to statistical analyses, the data were normalized by the median, log transformed using the base-10 logarithm (log10), and auto-scaled by mean-centering and dividing by the standard deviation of each metabolite feature to correct for non-normal distributions commonly observed in data^22^. Unsupervised analyses included Principal Component Analysis (PCA) and Hierarchical Cluster Analysis (HCA) to visualize overall metabolomic profiles. Additionally, supervised statistical analyses Partial Least Squares Discriminant Analysis (PLS-DA) and Orthogonal Partial Least Squares Discriminant Analysis (OPLS-DA) were used to visualize differences between SM and controls, as well as to identify specific populations of metabolites that discriminate between SM and control samples. In OPLS-DA, metabolite features are scored based on their discriminatory abilities using Variable Importance in Projection (VIP) scores. Metabolite features with the highest VIP scores have the greatest ability to discriminate between cohorts.

### Pathway analysis

Important metabolite features were selected using OPLS-DA VIP scores. Metabolites with the highest VIP scores (VIP > 1) from OPLS-DA were mapped to metabolic pathways using MetaboAnalyst’s Functional Analysis module with the *mummichog* algorithm which predicts a network of functional activity based on the projection of detected metabolite features onto local pathways^25^. Pathway library Human MFN was used for compound identification and pathway enrichment (mass tolerance: positive mode; 5 ppm; ranked by P-values; enforced primary ions (V2 only)). Pathways are reported as significant by pathway overrepresentation analysis with an adjusted *a priori* significance level of 0.1. The rationale for this significance level is that these are the first studies on the effects of microgravity on chondrocyte metabolomic profiles.

## Results

### Cell viability

A subset of SM and control constructs was analyzed for cell viability by staining with calcein AM and propidium iodide. The relative fluorescence was measured using a fluorescent plate reader. Cell viability assays confirmed that chondrocyte constructs were alive, with cell viability estimated at > 90% (Figure 2).

**Figure 2.**
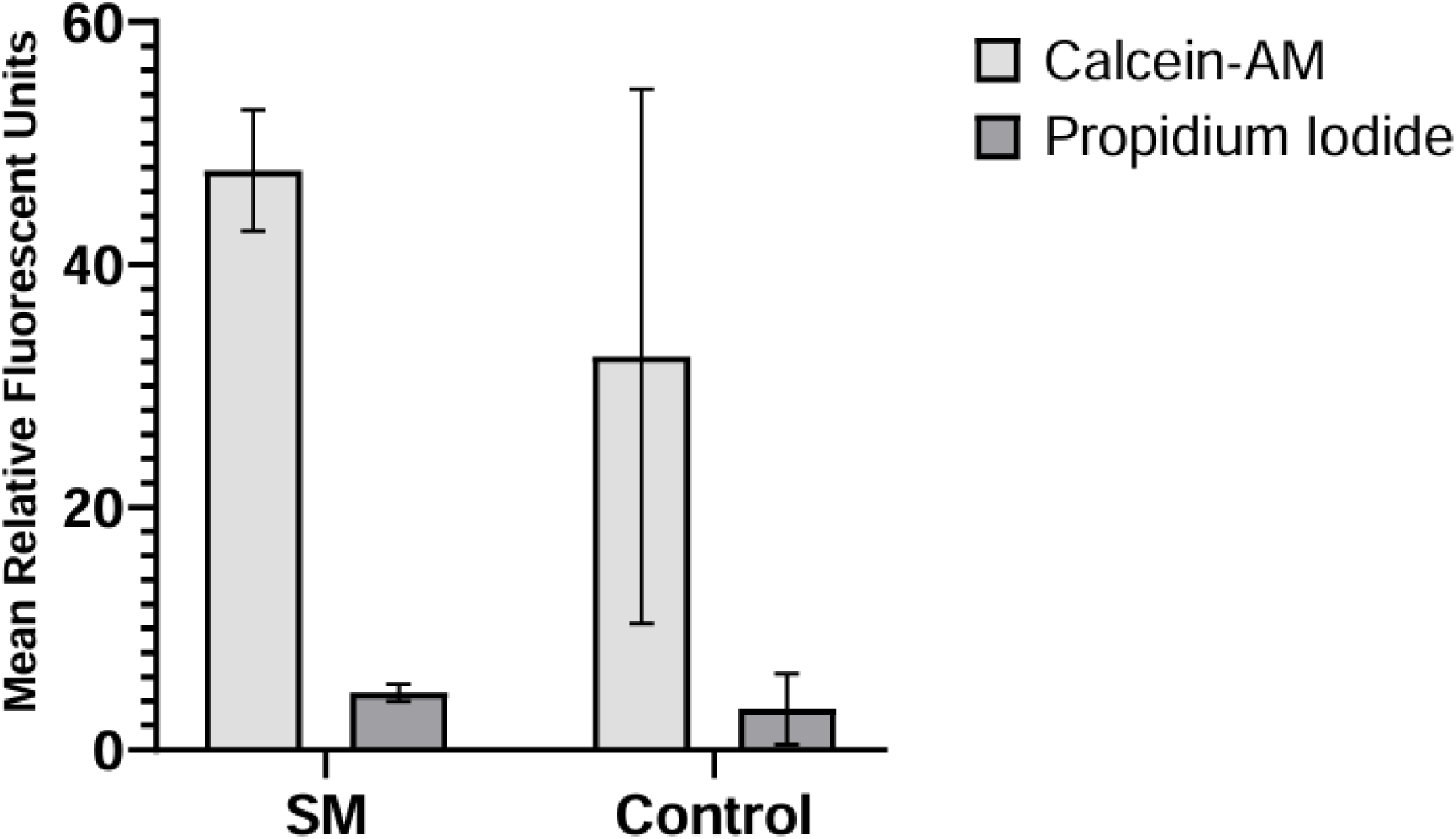
Viability of encapsulated chondrocytes. Mean relative fluorescent units for the SM (experimental) and normal gravity (control) cohorts with associated standard deviation bars.

### Global metabolomic profiling of constructs and media exposed to short-term simulated microgravity

Global metabolomic profiling detected a total of 1205 metabolite features across all samples, with 497 significant metabolite features identified by ANOVA (FDR-corrected p-value < 0.05). Unsupervised statistical analyses visualized the overall variability within the datasets and found minimal separation between control and SM cohorts in PCA plots and HCA dendrograms (Figure 3A-B). However, PCA demonstrated greater variability in the SM cohorts in comparison to the control cohorts, as exhibited by tighter clustering of control samples (Figure 3B). PLS-DA showed clear separation between gel and media cohorts, with some separation between control and SM cohorts (Figure 3C). These results suggest that short-term exposure may not induce large-scale shifts in overall metabolism (as shown by minimal separation in PCA); however, it is possible to tease out some metabolic shifts (as indicated by partial PLS-DA separation between control and SM cohorts).

**Figure 3.**
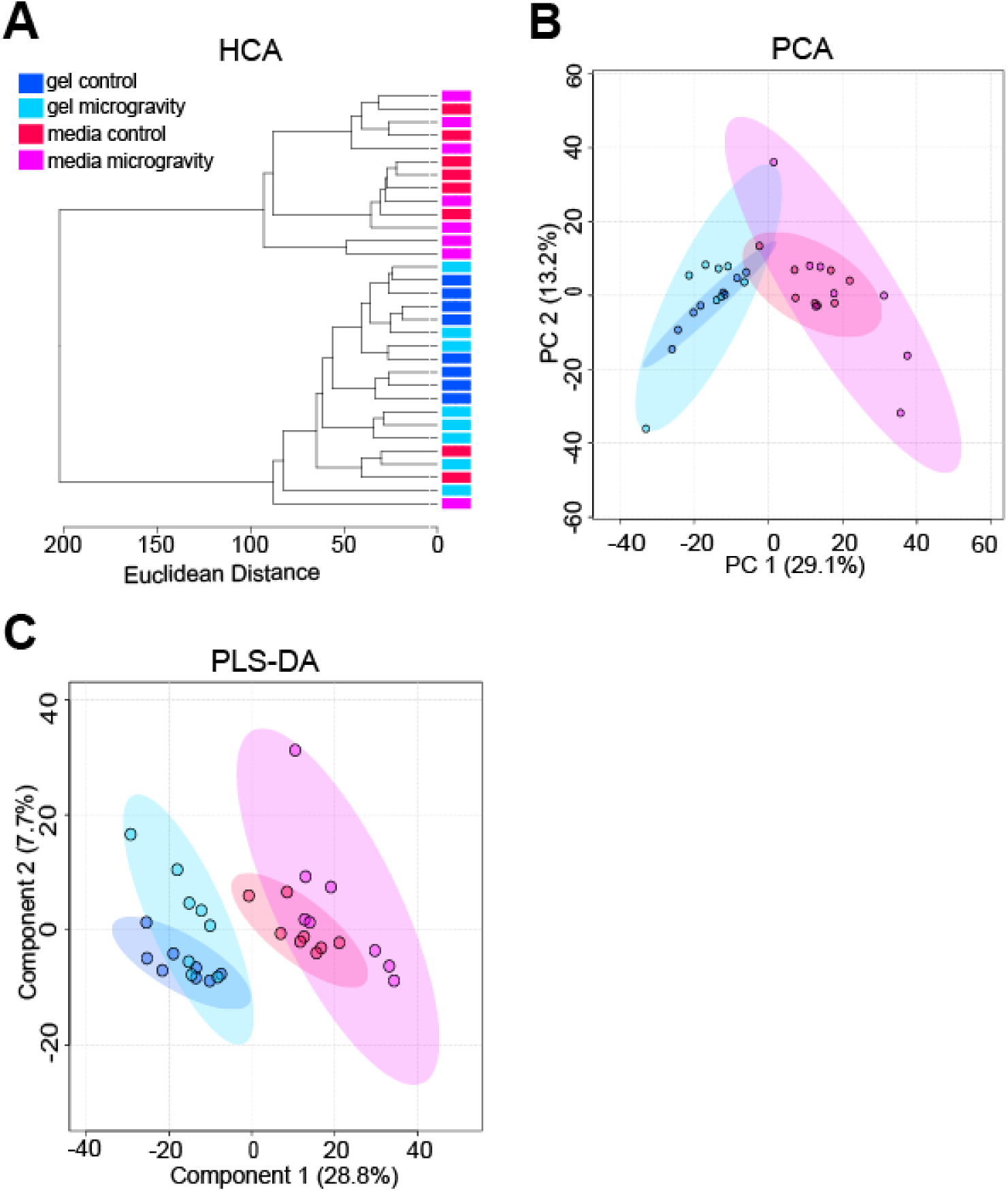
Unsupervised HCA and PCA, and supervised PLS-DA exhibiting clustering of constructs and media cohorts. **(A)** Unsupervised HCA dendrogram exhibiting separation between constructs and media, but minimal separation between SM and control samples. Clusters of cohorts were identified and labeled as gel control, gel microgravity, media control, and media microgravity. Each cohort contained 8 samples. Line length represents Euclidean distances between samples. **(B)** Unsupervised PCA plot exhibiting separation between SM and media, but minimal separation between SM and control samples. The two PCs are associated with 42.3% of the variation between cohorts (PC1 = 29.1%; PC2 = 13.2%). Samples correspond to their respective cohort: dark blue for gel control, light blue for gel microgravity, red for media control, and pink for media microgravity. **(C)** Supervised PLS-DA plot exhibiting separation between constructs and media, with some separation between SM and control samples. Component 1 and component 2 account for 36.5% of the overall variance (component 1 = 28.8%; component 2 = 7.7%). Samples correspond to their respective cohort: dark blue for gel control, light blue for gel microgravity, red for media control, and pink for media microgravity.

To pinpoint specific metabolic shifts associated with SM exposure, median intensity values were calculated for each metabolite feature. These medians were then clustered using HCA, and clusters were then subjected to pathway analysis (Figure 4) and enrichment (Table 1). Clusters of co-regulated metabolites were identified that represented microgravity-induced metabolic shifts: (1) differences between SM constructs and control constructs – clusters 3 (197 metabolites) and 5 (165 metabolites), and (2) differences between SM media and control media – clusters 1 (454 metabolites) and 4 (686 metabolites). Metabolites with higher abundances in encapsulated chondrocytes exposed to SM compared to control conditions mapped to amino sugar and nucleotide metabolism (Cluster 3, Figure 4, and Table 1). Conversely, metabolite features that were lower in abundance in SM media compared to controls mapped to glutathione, nitrogen, arginine, tyrosine, glutamine, glutamate, proline, phenylalanine, tryptophan, and aspartate metabolism (Cluster 1, Figure 4, and Table 1). Metabolite features lower in abundance in SM constructs compared to controls mapped to vitamin B_6_, tyrosine, porphyrin, and phenylalanine metabolism (Cluster 5, Figure 4, and Table 1). Metabolites higher in abundance in SM media compared to controls mapped to valine, leucine, isoleucine, glycine, serine, threonine, cysteine, methionine, and histidine metabolism (Cluster 4, Figure 4, and Table 1).

**Table 1.** Clustergram pathways. Pathway enrichment of HCA clustergram showcasing clusters of upregulated and downregulated metabolic pathways for gel vs. media differences. Clusters 1, 4, 3, and 5 correlate to HCA clustergram clusters in Figure 4. Clusters 2 and 6 from HCA clustergram did not reveal significant metabolic pathways. Pathway enrichment of HCA.

| **Cluster 1: ↑ Ctrl ↓ SM Media** | **Cluster 4: ↑ SM ↓ Ctrl Media** | **Cluster 3: ↑ SM ↓ Ctrl Constructs** | **Cluster 5: ↑ Ctrl ↓ SM Constructs** |
| --- | --- | --- | --- |
| Aminoacyl tRNA biosynthesis | Amino acid metabolism (valine, leucine, isoleucine degradation, metabolism of glycine, serine, threonine, histidine, cysteine, and methionine) | Amino sugar and nucleotide metabolism | Vitamin B<sub>6</sub> metabolism |
| Glutathione metabolism | Pantothenate and CoA biosynthesis | Valine, leucine, and isoleucine metabolism | Tyrosine metabolism |
| Amino acid metabolism (arginine, tyrosine, glutamine/glutamate, phenylalanine, tryptophan) | Propanoate metabolism |  | Porphyrin metabolism |
| Glyoxylate and dicarboxylate metabolism | Porphyrin metabolism |  | Phenylalanine metabolism |
| Pyrimidine metabolism | Aminoacyl-tRNA biosynthesis |  |  |
| Nitrogen metabolism |  |  |  |

**Figure 4.**
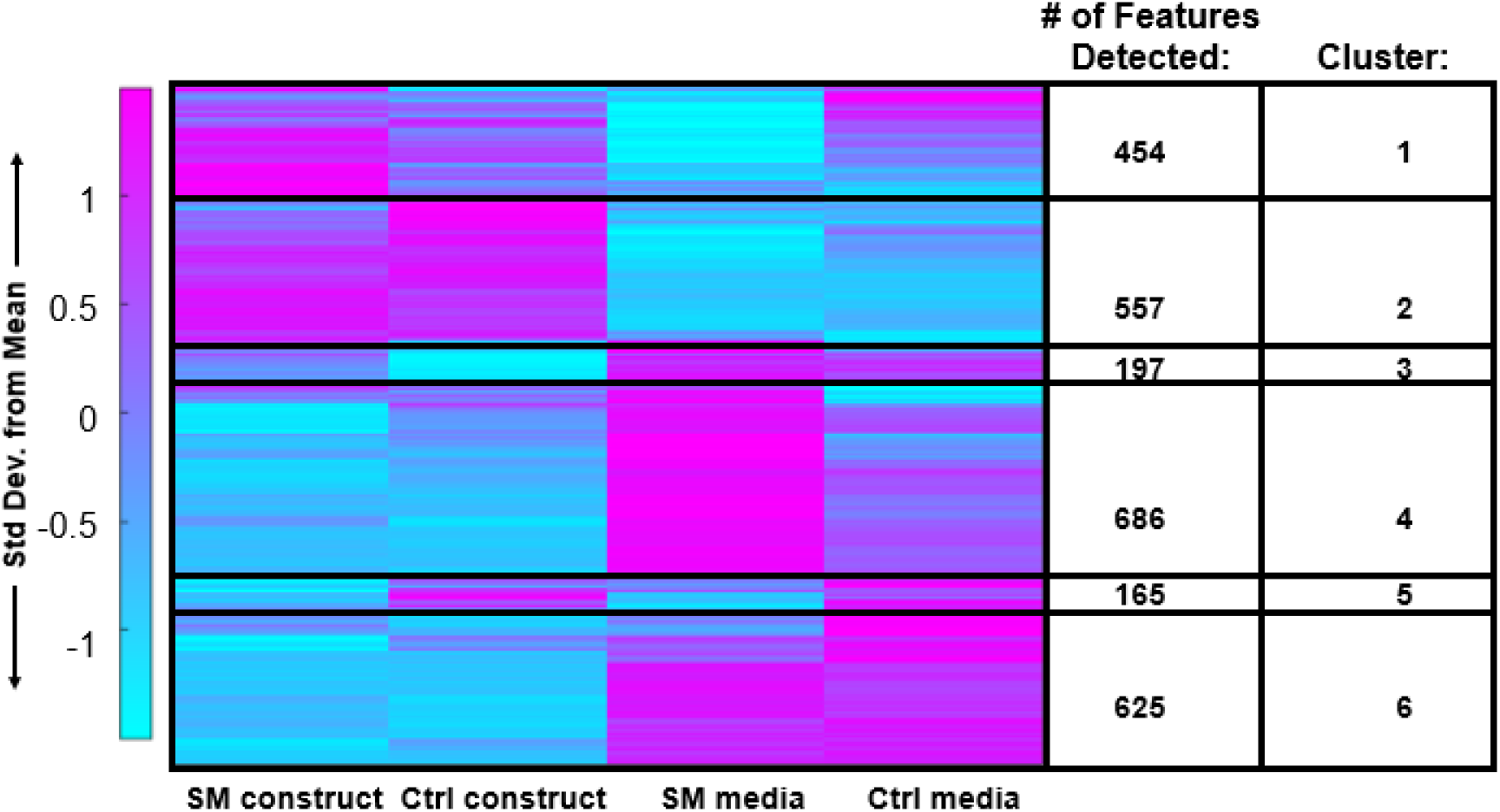
HCA clustergram showcasing global metabolomic profiles of simulated microgravity and control constructs and media. Rows represent metabolite features and columns represent experimental cohorts. The colors express if the specific metabolite feature was higher (magenta) or lower (blue) based on standard deviations from the mean metabolite intensity across all experimental cohorts. SM = simulated microgravity and Ctrl = control/normal gravity.

### Microgravity-induced metabolic shifts in chondrocyte constructs and simulated microgravity media

OPLS-DA was used to examine differences in metabolomic profiles between SM and control cohorts and identify specific metabolites features that contributed to separation between cohorts. OPLS-DA revealed clear separation between SM and control gel cohorts (Figure 5). Metabolites with the highest VIP scores contributed the most to the separation between SM and control gels and were mapped to metabolic pathways reported in Table 2. Select pathways perturbed in response to SM were mapped to di-unsaturated fatty acid β-oxidation, lysine, purine, vitamin B_6_ (pyridoxine), and urea cycle/amino group metabolism (FDR-corrected p-value < 0.1).

**Figure 5.**
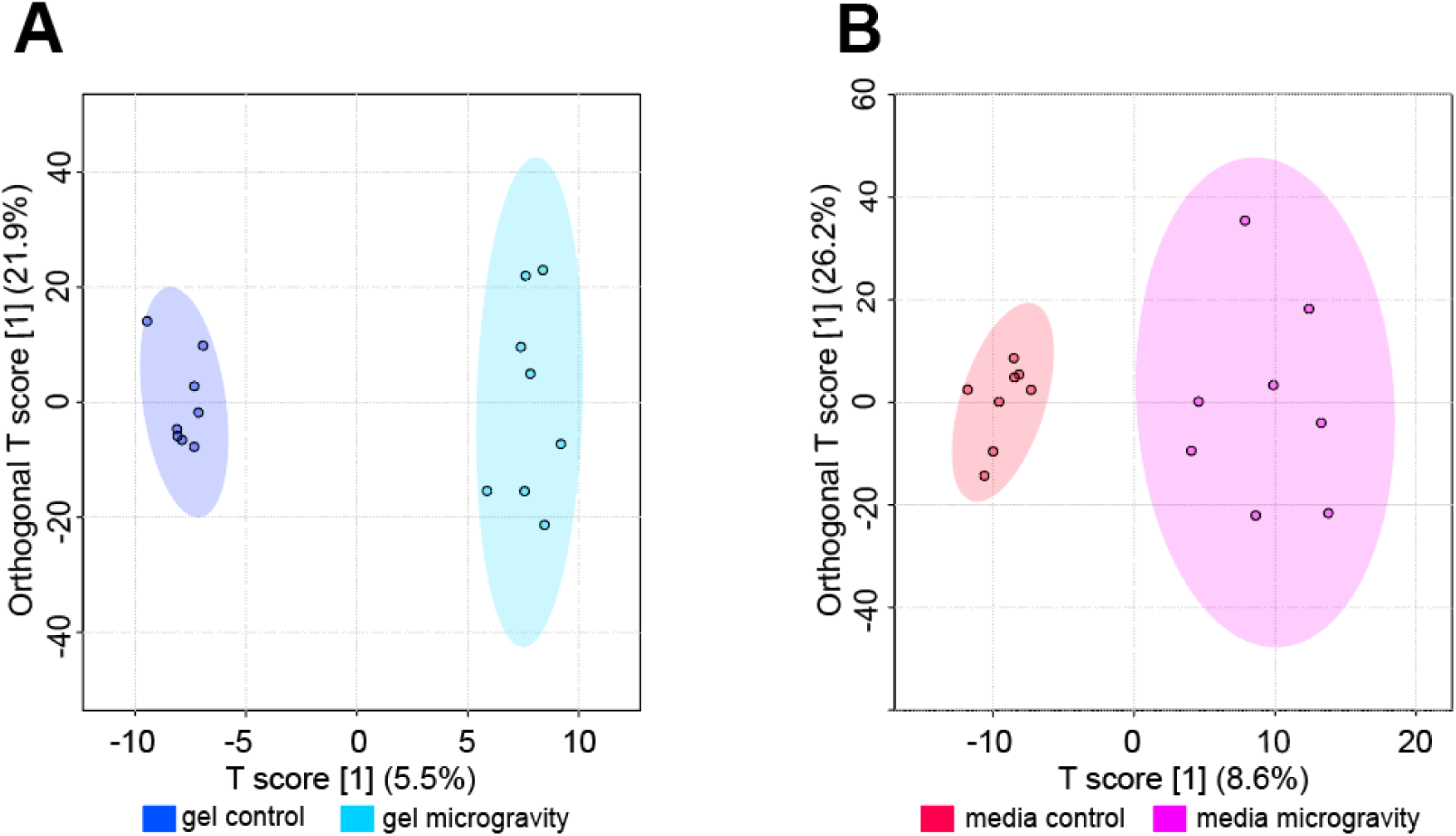
Supervised OPLS-DA plots exhibiting separation between control and SM samples. **(A)** Supervised OPLS-DA plot for gel samples exhibiting separation between control and SM samples. The T score showcases the inter-group variability (T score = 5.5%). The Orthogonal T score showcases the intra-group variability (Orthogonal T score = 21.9%). Samples correspond to their respective cohorts: dark blue for gel control and light blue for gel microgravity. **(B)** Supervised OPLS-DA plot for media samples exhibiting separation between control and SM samples. The T score showcases the inter-group variability (T score = 8.6%). The Orthogonal T score showcases the intra-group variability (Orthogonal T score = 26.2%). Samples correspond to their respective cohorts: red for media control and pink for media microgravity.

**Table 2.** OPLS-DA pathways – gels. SM altered metabolic pathways in gel constructs. Enrichment of gel SM and control samples mapped to significant metabolic pathways via OPLS-DA VIP Scores. The metabolites detected in this table are those of significance. The p-value reported is FDR-corrected. This is a condensed list of implicated pathways.

**Pathway Enrichment Analysis of OPLS-DA VIP for Gel Samples (VIP > 1)**
| Pathway | Metabolites Detected | Detected Metabolites with VIP $\geq 1$ | p-Value<sup>1</sup> |
| --- | --- | --- | --- |
| Di-unsaturated fatty acid β-oxidation | 4 | 3 | 0.026862 |
| Caffeine metabolism | 4 | 3 | 0.026862 |
| Xenobiotics metabolism | 4 | 3 | 0.026862 |
| Lysine metabolism | 16 | 6 | 0.032733 |
| Urea cycle/amino group metabolism | 25 | 8 | 0.037463 |
| Purine metabolism | 7 | 3 | 0.045989 |
| Vitamin B<sub>6</sub> (pyridoxine) metabolism | 3 | 2 | 0.047776 |
| Arginine and Proline metabolism | 16 | 5 | 0.050698 |
| Leukotriene metabolism | 26 | 7 | 0.059809 |
| Aspartate and asparagine metabolism | 28 | 7 | 0.07182 |
| Aminosugars metabolism | 5 | 2 | 0.07363 |
| β-Alanine metabolism | 5 | 2 | 0.07363 |
| Butanoate metabolism | 6 | 2 | 0.087549 |
| Omega-6 fatty acid metabolism | 6 | 2 | 0.087549 |
| Tryptophan metabolism | 17 | 4 | 0.093935 |
| <sup>1</sup> FDR-corrected |  |  |  |

OPLS-DA plot showed greater separation in SM media compared to control media (Figure 5). Metabolites with the highest VIP scores contributed the most to the separation between SM and control media and were mapped to metabolic pathways reported in Table 3. Select pathways perturbed in response to SM were mapped to glutathione, nitrogen, histidine, vitamin B_3_ (nicotinate and nicotinamide), and aminosugars metabolism (FDR-adjusted p-value < 0.1).

**Table 3.** OPLS-DA pathways - media. SM altered metabolic pathways in media. Enrichment of media SM and control samples mapped to significant metabolic pathways via OPLS-DA VIP Scores. The metabolites detected in this table are those of significance. The p-value reported is FDR-corrected. This is a condensed list of implicated pathways.

| Pathway Enrichment Analysis of OPLS-DA VIP for Media Samples (VIP > 1) |  |  |  |
| --- | --- | --- | --- |
| Pathway | Metabolites Detected | Detected Metabolites with VIP ≥ 1 | p-Value<sup>1</sup> |
| Glutathione Metabolism | 4 | 3 | 0.04789 |
| Glycosylphosphatidylinositol(GPI)-anchor biosynthesis | 2 | 2 | 0.059042 |
| Nitrogen metabolism | 2 | 2 | 0.059042 |
| Histidine metabolism | 10 | 5 | 0.062849 |
| Vitamin B<sub>3</sub> (nicotinate and nicotinamide) metabolism | 5 | 3 | 0.065442 |
| Amino sugars metabolism | 5 | 3 | 0.065442 |
| β-Alanine metabolism | 5 | 3 | 0.065442 |
| Androgen and estrogen biosynthesis and metabolism | 13 | 6 | 0.069171 |
| Glycerophospholipid metabolism | 16 | 7 | 0.075175 |
| Fatty Acid Metabolism | 6 | 3 | 0.085623 |
| Butanoate metabolism | 6 | 3 | 0.085623 |
| Phosphatidylinositol phosphate metabolism | 6 | 3 | 0.085623 |
| Methionine and cysteine metabolism | 6 | 2 | 0.085623 |
| Alkaloid biosynthesis II | 3 | 2 | 0.085726 |
| Vitamin E metabolism | 3 | 2 | 0.085726 |
| Vitamin A (retinol) metabolism | 3 | 2 | 0.085726 |
| C21-steroid hormone biosynthesis and metabolism | 17 | 7 | 0.88689 |
| <sup>1</sup> FDR-corrected |  |  |  |

## Discussion

### Overview

While recent research suggests that microgravity exposure leads to AC degeneration^10-17^, the underlying mechanisms driving this degeneration in reduced loading environments are currently unknown. Furthermore, there is a critical need for *in vitro* models that recapitulate the biomechanical microenvironment of chondrocytes for improved understanding of chondrocyte mechanotransduction in reduced loading environments. In the present study, we generated global metabolomic profiles of encapsulated chondrocytes exposed to SM to expand our understanding of chondrocyte mechanotransduction in reduced loading environments to better assess the risk of OA development post-spaceflight. To our knowledge, this is the first study to encapsulate chondrocytes in agarose gel constructs of similar stiffness to the PCM for suspension in a RWV bioreactor to examine the metabolic effects of short-term SM exposure.

### Mimicking the Chondrocyte Biomechanical Environment

A handful of *in vitro* studies document the effects of SM exposure on liquid cell culture of chondrocytes^26-30^. Experiments under SM are advantageous in that they circumvent many of the challenges of conducting experiments in space (*e.g.*, limited quantity, high cost). These studies suggest that chondrocytes exposed to SM exhibit cytoskeletal rearrangements, assembly of cells, and changes in gene expression^26-30^. While these studies provide insight into chondrocyte mechanotransduction in reduced loading environments, they do not consider the mechanical stiffness of the extracellular microenvironment and how it may impact mechanotransduction to chondrocytes.

Recapitulation of the chondrocyte pericellular microenvironment is necessary for studies of chondrocyte mechanotransduction^31,32^. It is well-established that chondrocytes are mechanosensitive cells that respond to mechanical stimuli by altering cellular metabolism and other processes to maintain homeostasis. *In vivo* chondrocytes are surrounded by a narrow (1-5 µm) layer of glycoproteins, proteoglycans, and collagen known as the pericellular matrix (PCM) that plays a key role in translating tissue level mechanical stimuli to chondrocytes. Importantly, there is a disparity in modulus between the PCM and surrounding extracellular matrix (ECM) in cartilage. Because the PCM is less stiff than the ECM, it can decrease the magnitude of stress applied to chondrocyts. Studies of chondrocyte mechanotransduction in response to increased physiological loading (*e.g.*, high impact loads) routinely embed chondrocytes in a 3D hydrogel to better mimic the cartilage microenvironment. Chondrocyte mechanotransduction studies in reduced loading environments, however, are limited to *in vitro* studies. This study takes the first step to improve *in vitro* modeling of chondrocyte mechanotransduction in reduced loading environments by embedding chondrocytes in an agarose gel of similar physiological stiffness to the PCM.

### Metabolic Response of Chondrocytes to Short-Term Simulated Microgravity

Current research suggests that deviations from physiological loading (*e.g.*, reduced or increased loading) result in an imbalance in chondrocyte metabolism, with catabolic processes predominating. Numerous studies characterize the metabolic response of chondrocytes to increased mechanical loads (*e.g.*, high impact loads)^2,3^. However, no study to date has investigated the metabolic effects of reduced loading on chondrocyte mechanotransduction.

The results of this study reveal that four-day exposure to reduced loading environments is sufficient to induce shifts in chondrocyte metabolism, which may contribute to the degeneration of AC. Exposure to SM induced significant alterations in amino acid metabolism and protein synthesis (valine, leucine, and isoleucine degradation, metabolism of glycine, serine, threonine, histidine, cysteine, methionine, arginine, tyrosine, phenylalanine, tryptophan, and vitamin B_6_, and aminoacyl tRNA biosynthesis). Additionally, significant changes in nucleotide metabolism (amino sugar and nucleotide metabolism, nitrogen metabolism, and pyrimidine metabolism), oxidative stress (glutathione metabolism), and energy metabolism (pantothenate and coenzyme A biosynthesis) were observed.

The metabolic response to short-term SM exposure suggests that chondrocytes respond to reduced loading by decreasing anabolic pathways required for ECM maintenance. This is shown by significant reductions in the metabolism of various amino acids and nucleotide metabolism in SM cohorts (constructs and media). While multiple studies show that chondrocytes respond to physiological loading by increasing the synthesis of ECM components^33-35^, the results herein suggest that reduced loading by SM leads to a reduction in the metabolic pathways necessary for the synthesis of ECM components, which may lead to inadequate maintenance of the cartilage structural integrity and degeneration consistent with that of OA.

Exposure to SM also induced shifts in chondrocyte energy metabolism, as shown by an increase in pantothenate and coenzyme A biosynthesis detected by metabolites secreted into surrounding media. Pantothenate (vitamin B_5_) is a precursor to coenzyme A, an important cofactor in the tricarboxylic acid (TCA) cycle. If not used to produce coenzyme A, pantothenate is exported from the cell. Increased levels of metabolites involved in pantothenate and coenzyme A biosynthesis detected in surrounding media may suggest that pantothenate is not being used for the production of coenzyme A, indicating that the energy requirements are lowered in response to the reduced mechanical loads. This is in line with a decrease in the anabolic pathways discussed above, which would significantly reduce chondrocyte energy requirements^36^.

Chondrocytes exposed to short-term SM exhibited a reduced ability to respond to oxidative stress, as shown by a decrease in the metabolism of glutathione, a known antioxidant. It is well established that microgravity exposure is associated with accumulation of reactive oxygen species (ROS) and a corresponding increase in oxidative stress. Increased oxidative stress alters chondrocyte metabolism, leading to an imbalance in matrix synthesis and degradation. Under normal conditions, chondrocytes are capable of counteracting oxidative stress through the production of antioxidants. However, the results herein suggest that even short-term exposure to SM may compromise the chondrocytes’ ability to counteract oxidative stress, which could lead to AC degeneration.

### Metabolic Pathway Alterations Consistent with Osteoarthritis

Many of the observed SM-induced metabolic perturbations have been previously implicated in OA. Several studies have shown differences in amino acid concentrations and amino acid metabolism in OA cartilage and biofluids (synovial fluid, urine, plasma), indicating their importance in OA pathogenesis(reviewed in 36). Additionally, OA chondrocytes exhibit alterations in energy metabolism, such as an increase in anaerobic glycolysis and reductions in the TCA cycle and oxidative phosphorylation^37,38^. Notably, a hallmark of OA is the chondrocyte’s inability to maintain homeostasis in response to oxidative stress^37, 39-44^. Studies suggest that glutathione levels are significantly reduced with aging and in OA^43, 44^. Taken together, these results suggest that even short-term exposure to reduced loading environments may induce early metabolic shifts consistent with that of OA.

Importantly, many similarities were observed between metabolites secreted from encapsulated chondrocytes exposed to short-term SM into the surrounding media and our previous work analyzing metabolomic profiles of OA synovial fluid^45^. Metabolites involved in the following pathways were perturbed in both SM media and early OA synovial fluid: histidine, amino sugars, fatty acid, butanoate, phosphatidyl phosphate, methionine, and cysteine metabolism, valine, leucine, and isoleucine degradation; glycine, serine, threonine, methionine, cysteine, glutathione, arginine, proline, and glutamate metabolism. These results provide further validation that reduced loading, even in the short-term, may induce metabolic shifts consistent with early OA. Furthermore, the detection of these metabolic perturbations as secretions into surrounding media suggest that microgravity-induced metabolic shifts may be detectable in human synovial fluid *in vivo* to assess osteoarthritis risk in astronauts.

### Limitations

This work has important limitations providing opportunities for future work. Future studies will increase the sample size of experimental cohorts to confirm microgravity-induced metabolic shifts detected in media and encapsulated chondrocytes and use targeted metabolomic profiling for confident identification of metabolite identities and implicated pathways. Furthermore, applying cyclical mechanical stimulation (e.g. compression) with and without simulated microgravity may provide further insight into the role of microgravity in chondrocyte mechanotransduction. Finally, this work is limited by the use of the SW1353 cell line, a chondrosarcoma cell line, that does not perfectly model the behavior of primary chondrocytes. Thus, primary chondrocytes obtained from human cartilage will be used for future work to model the *in vitro* microenvironment of cartilage more accurately.

### Significance and Conclusion

To our knowledge, this is the first study to generate global metabolomic profiles of 3D encapsulated chondrocytes in response to short-term SM exposure. Importantly, this work enhances our ability to study chondrocyte mechanotransduction in SM *in vitro* by encapsulating chondrocytes in a three-dimensional agarose construct of physiological stiffness to the PCM that better mimics the cartilage microenvironment. Metabolic pathways perturbed in response to SM were consistent with early OA, and thus better informs the health risks associated with space travel.

## Acknowledgements

The authors would like to acknowledge the Montana State University Mass Spectrometry CORE Facility for their assistance with mass spectrometry.

## Author Contributions

ARB assisted in designing experiments, performed metabolite extractions, ran LC-MS samples, analyzed data, and drafted the manuscript. MGG and EAH assisted in designing experiments, extracting metabolites, carried out cell viability assays, and assisted in analyzing data. EV and GC assisted in designing experiments. MC, KB, and PPB assisted in cell viability assays. HDW ran LC-MS samples and assisted in analyzing data. RKJ assisted in analyzing data. AKH designed experiments and analyzed data. All authors have read and revised the manuscript.

